# Xylazine and fentanyl co-administration delays wound healing in mice

**DOI:** 10.1101/2025.06.18.660449

**Authors:** Madigan L. Bedard, Roland F. Seim, Alex R. Brown, Calista A. Cline, Kristen N. Gilley, Luke A. Wykoff, Mitchell Huffstickler, Puja B. Nakkala, Shannon Rausser, Nabarun Dasgupta, Justin C. Strickland, Leon G. Coleman, Zoé A. McElligott

## Abstract

Xylazine, a veterinary sedative increasingly found in the unregulated drug supply, is associated with severe skin wounds in humans, particularly when co-used with fentanyl. Despite growing concern, the mechanisms underlying these wounds remain unclear. To investigate how xylazine and fentanyl affect wound healing, we administered subcutaneous injections of saline, xylazine (3.2 mg/kg), fentanyl (1.0 mg/kg), or their combination to female C57BL/6J mice for 28 days. After a standardized punch biopsy, wound closure was tracked for 14 days, with continued drug exposure. Mice receiving the xylazine-fentanyl combination exhibited significantly delayed wound healing compared to all other groups, as shown by slower closure rates and increased area under the healing curve. A follow-up study without chronic pretreatment showed that acute xylazine-fentanyl exposure still altered healing dynamics, although it did not significantly delay time to closure. Neither drug alone impaired healing at the tested doses. These findings suggest that prior exposure to xylazine and fentanyl contributes to impaired wound healing and support the hypothesis that xylazine-associated wounds may arise from delayed healing of pre-existing skin injuries rather than spontaneous formation. This is the first preclinical model of xylazine-related wound impairment and provides a foundation for future research into biological mechanisms and potential interventions for these emerging soft tissue injuries.

## Introduction

Xylazine, a common veterinary sedative, has been found in the unregulated drug supply at increasing rates over the last few years. This has posed significant challenges for medical and public health officials as it has drastically changed the withdrawal and treatment landscape. Xylazine exposure in humans is associated with necrotic skin wounds, increased sedation, and increased anxiety during withdrawal (1, 2). While xylazine use has been associated with wound development epidemiologically, we lack definitive evidence that wounds are causally related to xylazine as experimental studies in humans are unethical.

Initial reports suggested that wounds were “spontaneously forming” but more recent studies have found that wounds are developing where other injuries to the epidermis/dermis have occurred, often in conjunction with subcutaneous use (i.e., skin popping) (3). To our knowledge, no studies have reported spontaneous wound formation in animal models involving xylazine. We have previously investigated the impact of xylazine/fentanyl administration in C57BL/6J mice and reported sex-differences in withdrawal and locomotion responses over the course of multiple days and rounds of drug exposures(4). During the course of our experiments, we observed no spontaneous wound formation in male or female mice. Given the severity of wounds and lack of treatment options, we sought to develop an animal model to better understand the genesis of these wounds in people that use xylazine. Here, we present the first preclinical model of xylazine-impacted wound healing as a method for uncovering the underlying mechanism of wound formation and progression. Our data demonstrate that xylazine and fentanyl use impedes wound healing in female mice, suggesting this drug combination may contribute to the development of skin wounds in humans who use xylazine and fentanyl.

## Methods and Results

To assay wound healing, blinded investigators injected (s.c.) female C57BL6/J mice with saline, xylazine (3.2 mg/kg), fentanyl (1 mg/kg), or xylazine and fentanyl combined (3.2 and 1.0 mg/kg respectively, Extended Methods for additional dose discussion) for 28 days (Fig. 1A). We observed no spontaneous wound formation during the chronic drug exposure period. Following the 28-day pretreatment, animals were anesthetized (isoflurane) and wounds were made on the flank of the mouse (see Extended Methods and Materials) and measured using calipers (Fig.1B). During the wound healing assay that followed (14 days) mice were treated twice daily with the same drug regimen as during the pretreatment period. We observed a significant main effect of drug group and day post-injury on healing rate in a mixed-effects analysis (Fig. 1C; Drug: p<0.001, Time: p<0.001). There was also a significant interaction of drug and day (Fig.1C p<0.0001). To better assess how this impacted the healing, we completed two additional analyses: estimated time to heal with an independent regression analysis for each animal (Fig. 1D) and the area under the curve (AUC) of the group timelines (Fig. 1E). This shows the individual variability within each group. A one-way ANOVA revealed a significant main effect of drug group (p<0.0001, Fig. 1D), which was driven by the combination group that healed significantly slower than all other groups (Bonferroni’s post-hoc, Fig. 1D). The combination group had an effect size of 1.197 (Hedges’s *g*) compared to saline suggesting a large effect size (5). Additionally, we compared the area under the curve for the lines in panel C. This analysis provides insight into the pattern of healing as well as the rate of healing. The combination group had a significantly greater AUC than saline (Fig. 1E, p=0.004, Bonferroni’s post-hoc) and xylazine (p=0.0061, Bonferroni’s post-hoc), and was trending compared to fentanyl (p=0.1033). We found that the combination group had a Hedges’s *g* of 1.897 when compared to saline suggesting a large effect size of the treatment. Together, these results demonstrate the importance of the combination of fentanyl and xylazine in delaying wound healing, as neither drug alone significantly delayed healing.

**Figure 1.**
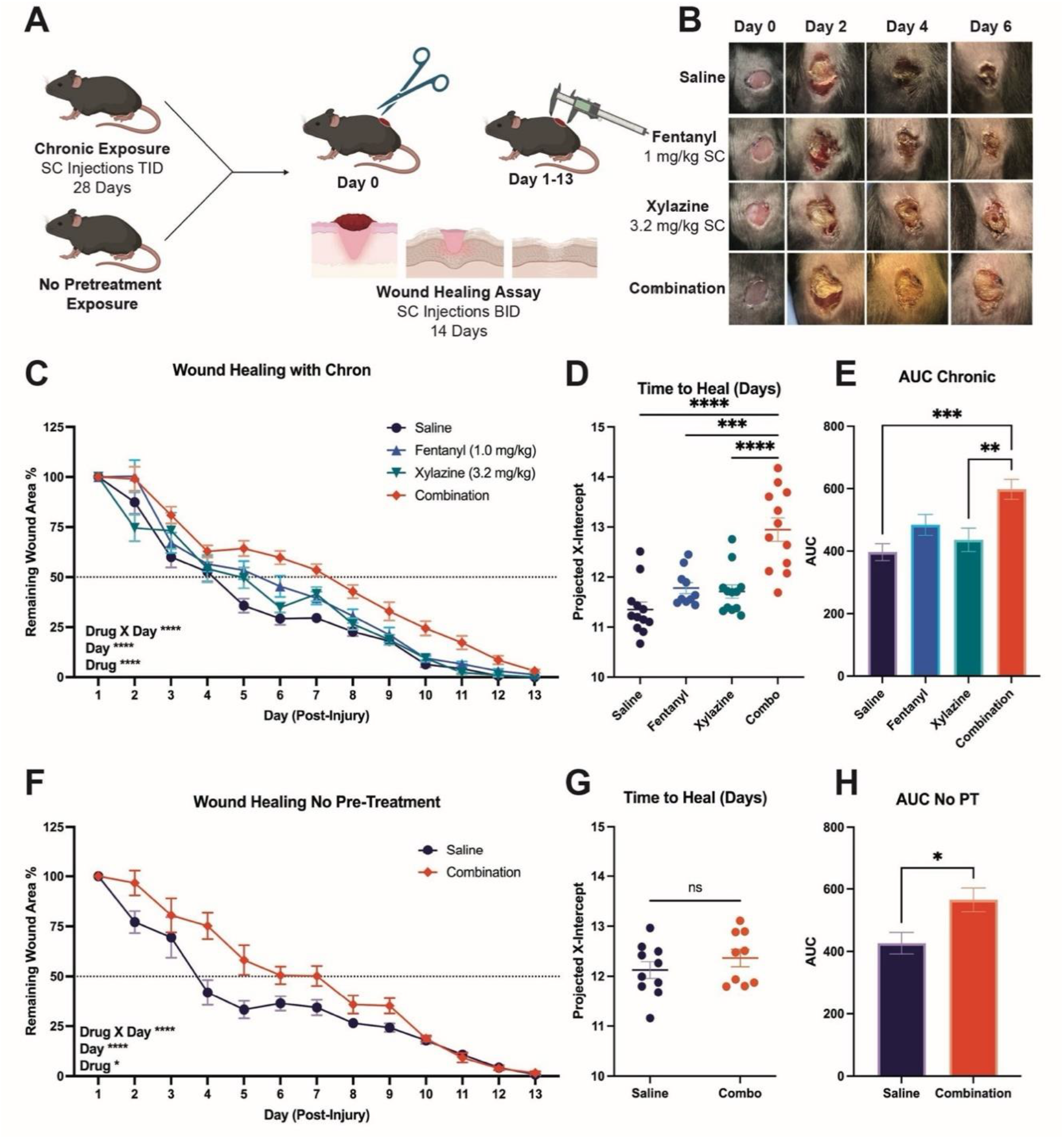
Xylazine and Fentanyl Delay Wound Healing. Paradigm for chronic and no pretreatment groups and wound healing assay (A). Representative images from each of the chronic exposure treatment groups during week 1 (B). Results of wound healing assay following chronic exposure as percentage of remaining wound area (C), time to completely heal in days (D), and area under the curve (E). Results of wound healing assay with no pre-treatment as percentage of remaining wound area (F), time to completely heal in days (G), and area under the curve (H). All percentages are relative to wound size on Day 1. p >0.05 ns, p ≤ 0.05 ^*^, p ≤ 0.01 ^**^, p ≤ 0.001 ^***^, p ≤ 0.0001 ^****^

Next, we repeated the wound healing assay in a new cohort of mice without the 28-day pretreatment. This allowed us to assess the importance of prior exposure to the drugs in wound healing. Because fentanyl and xylazine alone did not impact the overall wound healing rate in our chronic pretreatment study, we conducted this study with only the saline and combination groups. Again, we found that the drug group and day post-injury both significantly impacted wound healing and contributed to a significant interaction (Mixed-effects analysis, Drug: p=0.0126, Day: p<0.0001, Day X Drug p<0.0001, Fig. 1F). Without drug exposure prior to the wound, there was not a significant delay in the number of days it took the mice to fully heal (Fig. 1G); however, the combination treatment group did still have a significantly larger AUC, indicating a change in the pattern of healing (Fig. 1H, unpaired t-test p=0.0147). We found that the combination group had a Cohen’s *d* of 1.206 when compared to saline suggesting a large effect size.

## Discussion

### Prior Exposure to Xylazine + Fentanyl Impacts Wound Healing Rate

Our study finds that wound healing in female mice is negatively impacted by exposure to xylazine and fentanyl more than by either substance alone at the tested doses. Additionally, pre-exposure to the xylazine and fentanyl combination increased the time needed for wound closure. This work forms the basis for the first preclinical model of xylazine-associated wounds, a crucial step in our ability to determine the mechanistic underpinnings of this phenomenon. It remains to be determined if patterns of continuous exposure, alterations in dose and frequency of delivery, injection site location, or route of drug delivery would result in the same effect. We intentionally injected distally to the wound to observe how systemic administration would impact wound healing, however there are first person reports that individuals are injecting into the wounds directly (6). Evidence shows that most wounds are located on extensor surfaces of extremities, may be located at injection sites, or may be related to a non-injection route of administration such as inhalation/insufflation (1, 7). As venous access becomes difficult, people often search for new veins or routes of administration which may result in wounds in atypical locations, such as the skull (8).

Difficulty accessing veins might also increase the rate of subcutaneous injections (“skin popping”), increasing risk of wound development. Direct injection near painful wound sites may also become behaviorally relevant for localized anesthetic effects (9). One group reported a ten-fold increase in wound prevalence among those who injected subcutaneously compared to other routes (10).

Non-injection site wounds support the possibility that xylazine adulteration causes a delay in wound healing from an unrelated injury (e.g., bug bite or skin laceration) rather than causing the wound itself. Indeed, recent testimonies have suggested that wounds are appearing at the site of other insults to the epidermis (3). The lack of spontaneous wound formation in all animal studies also supports this theory, however, responses in humans and mice may differ and likely vary due to metabolic differences between species (11). Here, we show that female mice have significantly delayed wound healing in response to xylazine/fentanyl administration.

### Potential Mechanisms

While the mechanism of xylazine-impaired wound healing is still largely unclear, there are a few theories pertaining to the pharmacology of xylazine. Canonically xylazine is an α2-adrenergic receptor agonist. As such, it is possible that its effect on wounds is driven by α2-mediated vasoconstriction. Vasoconstriction leads to a reduction in angiogenesis, local ischemia, and limits the migration of cells to the wound resulting in impaired wound healing (12, 13). This factors may contribute mechanistically, since mice were ultimately able to close their wounds, albeit with significant delay.

We and others have recently established that xylazine is also an agonist at the kappa opioid receptor (кOR), sigma 1 and 2 receptors (σ1R and σ2R), as well as the serotonin (5-HT7A) receptor (4, 14). Each of these receptor systems provides an opportunity for interaction and could be evaluated for roles in wound formation and/or healing. Intriguingly, another кOR agonist, pentazocine, resulted in similar wound formation during the “Ts and Blues” outbreak from 1977-1981 (15).

Histologically, wound repair occurs in three stages: inflammation, new tissue formation, and remodeling (16). Combined treatment of fentanyl and xylazine may influence either some or all of these stages leading to a delay in wound closure. Immediately after injury, inflammatory pathways and parts of the coagulation cascade are activated to limit blood loss and to form a fibrin matrix over the wound. Next, neutrophils, monocytes, and keratinocytes are recruited to the site of the wound to coordinate repair, remove debris, and to form an eschar (scar) over the area. New blood vessels are formed and eventually the wound will consist primarily of collagen and extracellular matrix which is continuously remodeled over time. While the mechanism of wound formation in humans is unclear, our results showed that there is a delay in wound closure in mice that are treated with a combination of fentanyl and xylazine. Recent reports suggest that xylazine related wounds are developing at sites of prior injury (3).

Certain medications and/or other drugs can have a negative impact on wound healing rates by limiting cell proliferation, angiogenesis, cell migration, or immune function. Both µ-opioid receptor agonists and к-opioid receptor agonists have been shown to limit macrophage function by reducing their ability to produce pro-inflammatory cytokines (17, 18). Furthermore, кOR agonists may influence cellular and humoral immune responses on lymphocytes (19). To assess if chronic drug treatment had any effect on the peripheral immune populations, flow cytometry staining was performed on a subset of spleens from these mice at the conclusion of our study. In this preliminary study, we did not observe any changes in the overall number of peripheral immune populations of T-cells, monocytes, or myeloid-derived cells due to treatment. However, there could be deficiencies in immune cell function including recruitment of immune cells to the wound site, inhibited cytokine release, or limited phagocytic activity. Further experiments will need to be undertaken to rule out immune dysfunction as a primary mechanism for impaired wound healing following combined fentanyl and xylazine treatment.

An inability to adequately model wounds associated with xylazine use has left uncertainty regarding the mechanistic underpinnings of wound development and prolongation, and delayed identification of clinical targets. While mice have seven-fold faster metabolic rate as compared to humans (11), which undoubtedly affects the duration of exposure to the drugs, transgenic mouse models can afford unparalleled access to future mechanistic studies. This model enhances our ability to assess pathophysiology and potential therapeutics moving forward and can be expanded on as new or previously described substances are detected in the unregulated drug supply.

### Extended Materials and Methods Subjects and Drugs

All procedures described were approved by the University of North Carolina at Chapel Hill Institutional Animal Care and Use Committee (IACUC). Female C57BL/6J mice aged 11-15 weeks (Charles River Laboratories, Raleigh, NC) were maintained on a normal 12-hour light-dark cycle. All mice were group-housed and received food and water *ad libitum*. All drugs were delivered in 0.9% sterile saline solution and injected subcutaneously (SC) based on body weight. Xylazine injectable solution (3.2 mg/kg, Covetrus, Inc., Portland, ME) and fentanyl citrate (1.0 mg/kg, Spectrum Chemical MFG Corp, Gardena, California) were injected alone or in combination. Experimenters were blind to the administered drugs, and drug conditions were randomly assigned by cage to prevent stress transfer.

### Chronic Drug Exposure

Mice (11 weeks) were randomly assigned to receive saline (0.9%, N=12), xylazine injectable solution (3.2 mg/kg, N=12), fentanyl citrate (1.0 mg/kg, N=12), or fentanyl citrate + xylazine (N=13). The dose of xylazine was chosen based on reports of sedative response in people using xylazine (20). 3.2 mg/kg was a dose of xylazine we had previously published as sedative in female mice(4). Mice were injected 3 times daily (9 AM, 1 PM, 5PM) for 28 days. During this time, we monitored the mice for spontaneous wound formation. 4 days after the chronic exposure ended, mice began the wound healing assay.

### No Pretreatment Exposure

Mice were randomly assigned to receive saline (0.9%, N=10) or fentanyl citrate + xylazine (1.0 + 3.2 mg/kg, respectively; N=10). Mice habituated to the vivarium for one week prior to the start of the wound healing assay.

### Wound Healing Assay

Mice were anesthetized using isoflurane, and their backs shaved (21). A small wound (8 mm) was made with a sterile punch biopsy tool on the dorsal-caudal-lateral region. The size of the wound bed was measured with calipers daily for 2 weeks. Measurements were taken of the length and width of the wound, and an approximate area determined, areas are shown as % remaining wound area relative to Day 1 (first day post-op). Mice continued to receive 2 injections daily throughout wound healing (9 AM and 5 PM). Estimated time to heal was calculated by determining the equation of the line for each mouse’s individual healing rate and calculating the x-intercept (y=0% remaining wound area). All mice were 15 weeks old at the time of the wound assay, regardless of pretreatment/chronic exposure.

### Statistics

All statistics were calculated using GraphPad Prism version 10.4.1, one and two-way ANOVAs, mixed effect analysis, or t-tests were used to determine significance. Data is represented as mean ± SEM. Indications of statistical significance followed GraphPad p-value style (p > 0.05 ns, p ≤ 0.05 ^*^, p ≤ 0.01 ^**^, p ≤ 0.001 ^***^, p ≤ 0.0001 ^****^). All mice were included; none were determined to be outliers using a ROUT analysis.

## Acknowledgments

The chronic administration component of the project is supported by the Food and Drug Administration (FDA) of the U.S. Department of Health and Human Services (HHS) as part of a financial assistance award [Research Triangle Center of Excellence in Regulatory Science & Innovation, U01FD007857, PI: ZAM] totaling $1,453,116 from Center for Drug Evaluation and Research (CDER). The contents are those of the authors and do not necessarily represent the official views of, nor an endorsement, by FDA/HHS, or the U.S. Government.

The project was also supported by the National Institutes of Health (NIH) of the U.S. HHS. R01DA049261 (ZAM), F31DA056211 (MLB), T32GM135095 (ARB)

